# Selected clinical strains of *Mycobacterium avium* unlock *in vivo* models for NTM drug discovery programs

**DOI:** 10.64898/2026.07.14.738398

**Authors:** Tine Van Win, Wannes Brangers, Ellen De Pauw, Agustin Resendiz-Sharpe, Gert-Jan Wijnant, Kobe Bamps, Natalie Lorent, Greetje Vande Velde, Emmanuel André

## Abstract

*Mycobacterium avium* pulmonary disease is an emerging global health challenge for which drug development remains limited by preclinical models that rely on laboratory strains and invasive endpoint analyses. Here, we compared recent clinical *M. avium* isolates with the reference strain ATCC 700898 across macrophage, *Galleria mellonella*, and murine infection models and evaluated longitudinal micro-computed tomography (µCT) as a non-invasive tool to monitor disease progression and treatment response. While extracellular growth rates were comparable, clinical isolates demonstrated enhanced host-associated fitness and induced higher bacterial burdens and more severe pulmonary pathology in mice than the reference strain. These strain-dependent differences were detected by quantitative µCT imaging. Using the hypervirulent isolate MYC_0069, we further show that clarithromycin monotherapy and standard-of-care triple therapy significantly reduced bacterial burden and lung pathology. Together, these findings establish a clinically relevant chronic *M. avium* model that combines clinical isolates with longitudinal imaging to enable preclinical anti-mycobacterial drug evaluation *in vivo*.

## 1. Introduction

Non-tuberculous mycobacterial (NTM) pulmonary disease (PD) has emerged as a growing global health concern, driven predominantly by *Mycobacterium avium* in many countries (1–3). Although NTM-PD frequently affects immunocompromised individuals or patients with chronic lung diseases, an increasing number of cases occur in immunocompetent individuals, highlighting its increasing clinical significance (4). This expanding burden is accompanied by poor clinical outcomes, with five-year-all-cause mortality approaching 27% and increasing to approximately 50% in macrolide-resistant disease (5,6). The clinical burden is mirrored by substantial economic impact, with U.S. inpatient care costs alone estimated at $1.7 billion for approximately 181.000 cases in 2014 (7).

Treatment remains challenging. The current clinical standard-of-care for *M. avium* pulmonary disease consists of a prolonged combination regimen of at least three oral antibiotics; a macrolide (azithromycin or clarithromycin), a rifamycin (rifampin or rifabutin), and ethambutol, typically administered for a minimum of 6-12 months (8). Despite this multidrug approach, resistance is increasingly common (9). Clinical cure rates remain low, ranging from 32% to 62% (10). Only 21% of patients with macrolide-resistant disease achieve microbiological cure (4,6). Together, these trends highlight an urgent need for more effective therapeutic strategies against *M. avium* (6).

But, progress in treating *M. avium* disease is constrained by limitations of current preclinical models, particularly mouse studies, which are costly, ethically challenging, and heavily reliant on invasive endpoint analyses to evaluate disease progression and therapeutic efficacy. Among these, murine models remain the gold standard for studying *M. avium* pathogenesis and treatment response (11). However, these models rely heavily on invasive, single-endpoint readouts such as colony forming units (CFU) and histopathology (12). As a result, they are labor-intensive, time-consuming, and costly, while also raising ethical concerns due to the large number of animals required (13). Given the slow progression of NTM infection in humans and the prolonged duration of treatment, the lack of non-invasive, real-time readouts further limits their utility.

To address some of these limitations and facilitate early-stage virulence and drug screening, alternative infection models have gained increasing attention. In particular, the non-mammalian host *Galleria mellonella* has emerged as valuable intermediate platform between *in vitro* assays and mammalian studies. This model supports high-throughput screening, is cost-effective and ethically favorable, and can be maintained at 37°C, thereby allowing expression of temperature-regulated virulence traits (14). While its relative simplicity limits its scope, *Galleria* has proven useful for discriminating virulence differences among *M. avium* strains and may aid in identifying clinically relevant strain differences for downstream validation of mammalian models (15).

Within mouse infection models, there is a clear need for non-invasive longitudinal tools that enable real-time monitoring of disease dynamics within the same animal, reducing variability and animal use (14,16). Micro-computed tomography (µCT) offers such an opportunity by allowing real-time visualization and quantification of pulmonary pathology *in vivo.* By enabling automated and quantitative assessment of lung lesions, µCT overcomes key limitations of conventional endpoint-based analyses. Importantly, this approach facilitates longitudinal tracking of infection-associated lung changes over extended periods, enabling assessment of disease progression and treatment response within the same animal. The technique has been validated in multiple pulmonary disease models, including cryptococcosis, influenza-associated pulmonary aspergillosis, COVID-19, and bleomycin-induced lung fibrosis (17). Integrating µCT into *M. avium* infection models increases sensitivity for detecting early and strain-dependent differences in disease progression and provides a refined platform for evaluating treatment responses while reducing animal use. Overall, µCT enables dynamic monitoring of lung pathology, addressing the static and invasive nature of traditional histology and CFU-based assessments (2,18).

Beyond these technical and methodological limitations, a more fundamental challenge lies in the widespread use of laboratory reference strains, which fail to capture the extensive genetic, phenotypic, and virulence diversity observed among clinical isolates (3). As a result, even well-optimized experimental models may fail to accurately predict clinical outcomes when they rely on non-representative bacterial strains. For example, the widely used *M. avium* Chester strain (ATCC 700898), isolated in 1983, may not reflect contemporary virulence characteristics or emerging virulence trends (20). In contrast, clinical strains often display enhanced immune evasion and altered antibiotic tolerance, which can lead to underestimation of pathogenic potential and reduced predictive value in drug efficacy studies (3,7,8).

In this study, we compare selected clinical *M. avium* isolates with a commonly used reference strain to assess differences in virulence and disease progression across multiple infection models. To this end, we evaluate whether *Galleria mellonella* can serve as a predictive intermediate system capable of capturing strain-specific virulence traits, and determine whether micro-computed tomography (µCT) enables sensitive, non-invasive, and longitudinal detection of strain-specific differences in pulmonary pathology and treatment efficacy in mice. Together, this integrated approach demonstrates how *M. avium* clinical isolates can expose fundamental shortcomings of widely used experimental systems and provides a framework for the development of more predictive and clinically relevant models for NTM drug discovery.

## 2. Materials & methods

### 2.1 Bacterial strains and culture conditions

To assess differences in virulence among *Mycobacterium avium* strains, three strains were included in this study: the reference *M. avium* ATCC 700898 strain and two *M. avium* clinical isolates, internally pseudonymized as MYC_0068 and MYC_0069. Clinical isolates were selected from a collection of recent respiratory samples (isolation year 2020 and 2023, respectively) of patients meeting the clinical, microbiological and radiological patterns of pulmonary MAC disease, under ethical approval from UZ Leuven (approval number S72277). Species identification was confirmed by in-house whole genome sequencing. MIC testing all strains was performed to confirm similar baseline clarithromycin, ethambutol and rifampicin susceptibility.

All strains were stored in 25% glycerol (VWR, USA) at −80 °C until use. After thawing, all strains were grown on solid Middlebrook 7H10 agar medium containing 0.5% glycerol and 10% Oleic acid-Albumin-Dextrose-Catalase (OADC) (Fisher Scientific, USA) and incubated at 37°C for 10-14 days. Once a single CFU had grown sufficiently, it was transferred to Middlebrook 7H9 broth supplemented with 0.5% glycerol, 0.05% Tween- 80 (Merck, USA) and 10% Albumin-Dextrose-Catalase (ADC) (Merck, USA). Liquid cultures were incubated at 37°C, while shaking at 150 rpm. Bacterial concentrations in the exponential growth phase were estimated based on an optical density (OD_600_) to CFU correlation (CFU/mL) using a spectrophotometer (VWR, USA). Suspensions were centrifuged, resolubilized in PBS and subsequently diluted in PBS to the desired concentration. The final bacterial load was confirmed by CFU plating.

### 2.2 *In vitro* characterization of reference and clinical *M. avium* strains

*In vitro* infection experiments, including extracellular growth, intracellular macrophage infection, and infection rate were assessed for the reference strain and clinical isolates.

Extracellular growth kinetics were evaluated by culturing strains in Middlebrook 7H9 broth supplemented with ADC at 37 °C and was confirmed by CFU enumeration at predefined time points (see section 2.5).

For intracellular infection assays, THP-1 derived macrophages were seeded at 5×10^5^ cells/well in a 96-well plate, differentiated into macrophages using RPMI (Fisher Scientific, USA) + 10% FBS medium containing 500 ng/mL PMA (Merck, USA) over 3 days. On the day of infection, 5×10^5^ macrophages were infected with *M. avium* strains at a defined multiplicity of infection (MOI) of 1 for 4 hours. After phagocytosis, extracellular bacteria were removed by washing trice with PBS and replaced with RPMI + 10% FBS (Life technologies, USA). Infected macrophages were incubated at 37°C, 5% CO_2_ for up to 7 days, and at selected timepoints intracellular bacterial burden was quantified at by CFU enumeration following macrophage detachment by removing medium and adding 5mM EDTA in PBS. Afterwards macrophages are lysed using a 1/1 ratio of lysis buffer 0,1% triton in PBS for 5 minutes. All experiments were performed in biological triplicates. Medium was refreshed on day 3 to maintain host cell viability.

### 2.3 Establishment of infection models for comparative virulence assessment

#### 2.3.1 *Galleria mellonella* infection model

Bacterial inocula of the reference strain and the two selected clinical strains were used to infect healthy, 6^th^ instar *Galleria mellonella* larvae (in-house bred) weighing 300 ± 50 mg, with normal movement and no melanization. Larvae were randomly assigned to experimental groups (n = 12 per group) and housed individually in 12-well transparent plates (Greiner, Austria) at 37°C in the dark without food to mimic human host temperature.

To assess virulence differences, bacterial suspensions (10 µL PBS containing 10^6^ CFU) of the reference strain *M. avium* ATCC 700898 (16) and *M. avium* clinical isolates MYC_0068 and MYC_0069 were injected into the last left proleg using a Hamilton syringe (25 µL/ga30/12.7 mm Hamilton, USA). Negative controls were sham-infected with 10 µL PBS. At defined timepoints, larvae were homogenized and bacterial burden was quantified by CFU enumeration as described in section 2.5.

#### 2.3.2 BALB/c murine infection model and imaging

Female BALB/c mice (6-8 weeks old; Charles River Laboratories, Germany) were acclimatized for seven days in a biosafety level 2 facility prior to infection. Mice were housed in individually ventilated cages with free access to food and water. All murine experiments in this study were approved by the Ethics Committee on Animal research at KU Leuven (ECD license P148/2024).

Mice (n = 36) were randomized into groups of four per cage and intranasally infected with 25 µL of *M. avium* strains ATCC 700898 (2.5×10^5^ and 2.5×10^6^ CFU), MYC_0068 (2.5×10^5^ CFU), and MYC_0069 (2.5×10^5^ CFU). Anesthesia induction for inoculation was performed by gas anesthesia with 2% isoflurane (Piramal Healthcare, UK) in 100% O_2_ (nasal cone).

At experimental endpoints (weeks 4, 10, 16, and 26), samples (right lung, spleen, and liver) were collected and weighed. Left lungs were fixed for histopathological analysis (hematoxylin and eosin (H&E) and Ziehl-Neelsen staining). Lung lesion distribution was visualized using heat maps generated in the open-source QuPath software (version 0.5.1), applying a density radius of 200 µm, the Jet (legacy) colormap, a value range of 0-1502, an opacity range of 0-1502, and gamma = 1. Macroscopic images of the lungs were captured to support comparative pathology assessments.

Longitudinal monitoring included twice-weekly clinical scoring (body weight, respiratory parameters, general condition) and weekly µCT to assess lung pathology. µCT data were acquired using a small animal µCT scanner (SkyScan 1278, Bruker micro-CT, Belgium) and the following scan parameters: 50 kV X-ray source, 1 mm aluminum X-ray filter, 350 µA current, 150 ms exposure time per projection, with three projections acquired and averaged, 0.9° rotation increments, partial width of 50%, and isotropic voxel size of ∼ 50 µm (21). Each animal was anesthetized using a gas mixture of 2-3% isoflurane in 100% O_2_ and a flow of 1L/min. After scanning, mice are placed on a heating pad to recover. Respiratory weighted µCT-data were reconstructed and visualized using NRecon (version 1.7.5.9) and Data Viewer (version 1.6.0.6) software provided by the manufacturer. The following parameters were used for image reconstruction: smoothing 2 and beam hardening correction of 10%; ring artefact reduction and post alignment were adjusted per scan. A µCT derived biomarker for lung pathology (non-aerated lung volume (NALV)) was subsequently quantified, using a generic deep learning-based lung segmentation model (17). Sacrifice was performed at predefined endpoints or if humane endpoints were reached using pentobarbital overdose. A schematic overview of the experimental design is shown in **Supplementary Figure 1A**.

### 2.4 Evaluation of treatment response using *M. avium* strains

#### 2.4.1 *Galleria mellonella* infection model

Based on the comparative virulence assessment results of the two clinical isolates (MYC_0068 and MYC_0069) in *Galleria* and mice, the most virulent isolate (MYC_0069) was selected for further treatment response testing in both *in vivo* models.

Larvae were infected with either the reference *M. avium* strain ATCC 700898 or the *M. avium* clinical isolate MYC_0069. Clarithromycin (CLA; VWR, USA, Catalog Nr. TCIAC2220) solution was diluted in PBS containing 0.6% acetic acid (Thermo Scientific, USA) to obtain final doses of 100, 10 and 1 mg/kg. Based on an average larval weight of 300 mg, treated and sham-treated larvae received a total injection volume of 10 µL of either clarithromycin or vehicle (0.6% acetic acid in PBS). Treatments were freshly prepared and administered by injection into the last proleg, starting from one day post infection and repeated to seven days post infection. Treatment response was assessed using the same read-out parameters as described for the *Galleria* infection model (section 2.3.1), including CFU enumeration from larval homogenates (see section 2.5).

#### 2.4.2 BALB/c murine infection model and imaging

To assess treatment efficacy and determine whether µCT sensitively detects therapeutic responses, female BALB/c mice (6-8 weeks old; n = 12 per group; Charles River Laboratories) were randomly assigned to four groups. Groups 1, 2, and 3 were intranasally infected with 25 µL *M. avium* strain MYC_0069 suspension (2.5×10^5^ CFU). Anesthesia induction for inoculation was performed by gas anesthesia with 2% isoflurane in 100% O_2_ (nasal cone). Mice received antibiotic or vehicle treatments: group 1 received clarithromycin, group 2 received a triple combination of clarithromycin (100 mg/kg), ethambutol (ETH; Merck, USA Catalog Nr. E4630-25G; 100 mg/kg), and rifampicin (RIF; Fisher Scientific, USA, Catalog Nr. 15483539; 20 mg/kg). Group 3 received vehicle only (0.6% acetic acid in PBS). Group 4 was uninfected (negative control) and received vehicle only (22). Treatments were administered once daily, seven days per week, for up to 12 weeks via oral gavage using a 20-gauge plastic feeding tube (Instech, USA) attached to a 1 mL sterile syringe (Henke-Ject, catalog no. HS 8300063458). Formulations were prepared weekly under sterile conditions, stored at 4°C, and protected from light.

As described in 2.3.2, right lung, spleen, and liver were collected and analyzed to assess mycobacterial burden at day 1, and week 4, 10 and 16 post infection. Bacterial counts were determined by CFU plating as described below. Left lungs were fixed for histopathological analysis (hematoxylin and eosin (H&E) and Ziehl-Neelsen staining). Lung lesion distribution was visualized as described before. Macroscopic images of the lungs were captured to support comparative pathology assessments. Likewise, longitudinal follow-up included twice-weekly clinical scoring (body weight, respiratory parameters, general condition) and weekly µCT to assess lung pathology as described in 2.3.2. The experimental design is shown in **Supplementary Figure 1B**.

### 2.5 Colony forming units (CFU)

To monitor mycobacterial burden, CFU counts were determined from larval, lung, spleen, and liver homogenates. Samples were homogenized in 1 mL 7H9 medium using a tissue homogenizer (Tissue Master homogenizer; Omni International, USA). Ten-fold serial dilutions of both the inocula (to verify the administered bacterial dose) and homogenates were prepared. 25 µL from each dilution were plated on 7H10 agar plates, incubated at 37°C, and colonies were counted after 14 days to determine CFU/mL for the larval homogenates and CFU/gram for the right lung, liver and spleen homogenates. Activated charcoal (0.4%) (Merck, USA) was added to the agar to inactivate any residual drug compounds at week 16.

### 2.6 Statistical analysis

All statistical analyses were performed using GraphPad Prism version 8.0.2. Log_10_-transformed bacterial burden (CFU/gram), NALV, mouse weight and weight of the right lung, liver and spleen were analyzed by two-way ANOVA (or mixed model) to test for significant differentiation between strains. The results were considered statistically significant when p < 0.05.

## 3. Results

### 3.1 *In vitro* characteristics differ between reference and clinical *M. avium* strains

Virulence differences among 3 tested *M. avium* strains (two clinical isolates and one reference strain) were assessed across 3 distinct infection models of increasing complexity and biological relevance: *in vitro* human macrophages, *in vivo* in *Galleria mellonella* larvae and in BALB/c mice.

In the simplest model, extracellular growth kinetics and intracellular macrophage infection were assessed. All strains demonstrated comparable extracellular growth rates under standard culture conditions (**Figure 1A**). Differences emerged when host-associated infection models were employed. In the macrophage infection assay, the clinical isolates showed enhanced mycobacterial growth compared with the reference strain (**Figure 1B**). Also the infection rates tended to be higher for the clinical isolates than for the reference strain, with MYC_0069 and MYC_0069 exhibiting infection rates of 1.05% and 4.13%, respectively, compared with 0.71% for ATCC 700898 (**Supplementary figure 2**). Although these differences were not statistically significant, the increased infection rates and intracellular persistence suggested a more virulent phenotype for MYC_0068 and MYC_0069. These isolates were therefore selected for further evaluation in *Galleria mellonella* and murine infection models.

**Figure 1.**
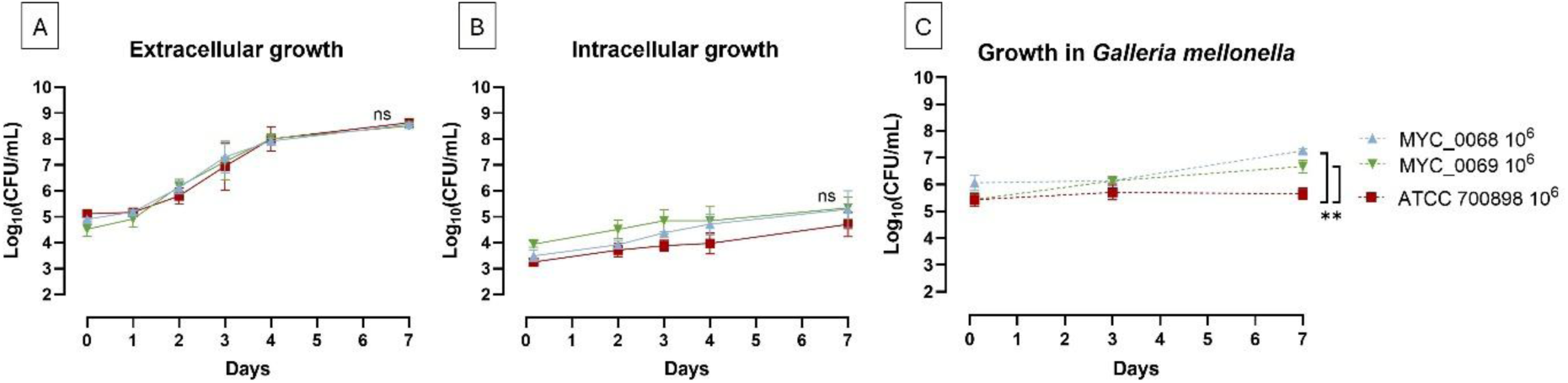
Clinical isolates exhibit enhanced host-associated growth. **A)** Extracellular growth kinetics of *M. avium* ATCC 700898, MYC_0068, and MYC_0069 under standard culture condition, monitored over time. **B)** Intracellular bacterial burden in macrophages following infection with the reference strain and clinical isolates, quantified at indicated timepoints post infection. **C)** Larvae were infected with 10^6^ CFU/10 µL of *M. avium* ATCC 700898, MYC_0068, or MYC_0069. Bacterial burden (log_10_(CFU/mL)) was quantified at 1 hour, 3 days, and 7 days post infection. Bars indicate ± SD (n = 4 per timepoint). Statistical significance: **P < 0.01, ns = non-significant.

### 3.2 Clinical isolates result in higher bacterial burden than the reference strain in *Galleria mellonella*

To determine whether virulence differences between clinical isolates and the reference strain were maintained in an invertebrate model, *G. mellonella* larvae were infected with 10^6^ CFU/10 µL of the three *M. avium* strains (**Figure 1C**). From 1 hour to 3 days post-infection, bacterial burden increased most for clinical isolate MYC_0069, while MYC_0068 and the reference ATCC700898 strain showed a comparable, stable bacterial burden. Between days 3 and 7, the reference strain again showed minimal replication, with no notable increase in CFU. In contrast, both clinical isolates demonstrated active bacterial growth over the same period, resulting in significantly higher bacterial burdens at day 7. This data confirms that the selected clinical isolates replicated more robustly than the reference strain in *G. mellonella*, with the growth rate varying between clinical isolates.

### 3.3 Clinical isolates establish stronger and more persistent lung infection in mice than the reference strain

To extend the virulence comparison to a mammalian host, BALB/c mice were intranasally infected with two different inocula of the reference ATCC 700898 strain (2.5×10^5^ and 2.5×10^6^ CFU/25 µL) or with an equivalent inoculum (2.5×10^5^ CFU/25 µL) of each clinical isolate. All strains transitioned from an acute to chronic infection phase. However, bacterial loads differed markedly between strain types (**Figure 2A**). Both clinical isolates maintained bacterial burden approximately 3.5log_10_ higher than the reference strain given at the same inoculum and at a ten-fold higher inoculum (2.5×10^6^ CFU) throughout the entire 26-week infection period. Although bacterial burden immediately after inoculation was not determined, comparison of MYC_0069 inocula across independent experiments indicated comparable infectivity (**Supplementary figure 3**). A similar trend regarding bacterial burden was confirmed in the spleen and liver (**Supplementary figure 4A-B**). These data further support that clinical isolates replicate substantially more robustly than the laboratory reference strain *in vivo*.

**Figure 2.**
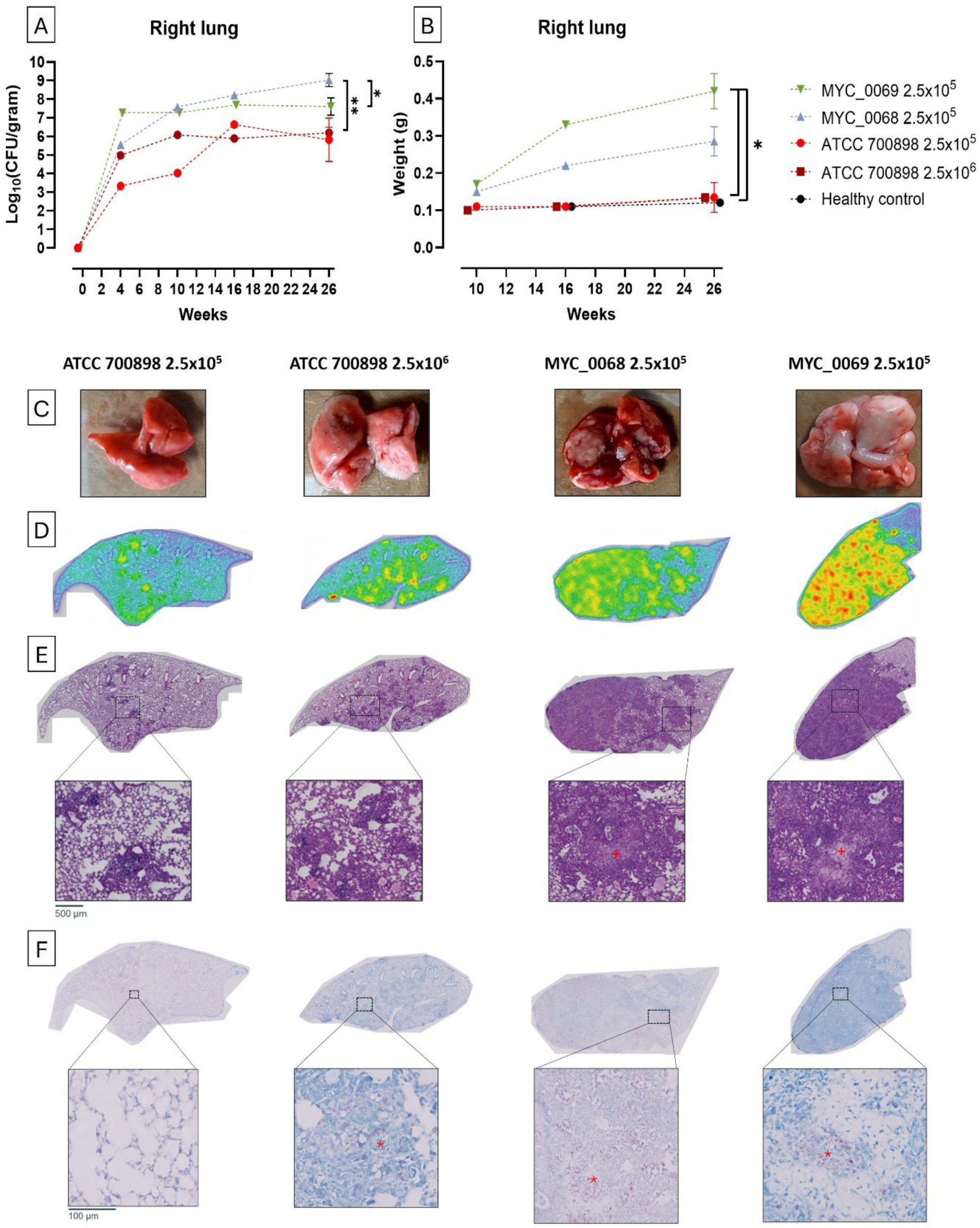
Clinical isolates cause more severe macroscopic and microscopic lung pathology than reference strain. BALB/c mice were intranasally infected with 2.5×10^6^ CFU of *M. avium* ATCC 700898 or 2.5×10^5^ CFU of *M. avium* ATCC700898, MYC_0068, or MYC_0069. **A)** Lung bacterial burden over time, expressed as log_10_(CFU/gram), measured at weeks 4, 10, 16, and 26 post infection. **B)** Lung weights at corresponding timepoints. Lung weights at week 4 are not shown because residual BAL fluid increased lung weight, which was avoided at later timepoints. Bars indicate mean ± SD for each group (n = 1/2 per infected/healthy group at week 4, n = 1 per group at week 10 and 16; n = 5/1 per infected/healthy group at week 26). **C)** Representative images of the right lung from mice infected with different *M. avium* strains and inoculum sizes at week 26; intermediate timepoints are not shown. **D)** Each H&E image was converted to heat maps. Red represents the lesion areas, identified by the dense presence of nuclei, green the thickening of parenchyma, and blue the uninvolved parenchymal tissue. **E)** H&E staining of the left lung showing whole-lung overviews and magnified regions (scale bar: 500 µm). Granuloma-like structures are indicated with the symbol (+). **F)** Ziehl-Neelsen staining of the same lung sections shown in (B), with magnified views highlighting acid-fast bacilli (scale bar: 100 µm). Acid-fast bacilli are indicated with the symbol (*). Statistical significance: *P < 0.05, **P < 0.01.

Neither inoculum of the reference strain increased lung weight relative to uninfected controls, suggesting minimal pulmonary pathology (**Figure 2B**). In contrast, both clinical isolates induced increased lung weights over time, with MYC_0069 showing significant enlargement compared with both reference strain inocula (**Figure 2B**). A similar effect was seen on the weight of the spleen but not of the liver (**Supplementary figure 4C-D**).

To assess the impact of *M. avium* infection on lung structure, lungs were imaged at each timepoint and histopathological analyses were performed. Marked structural differences between lungs infected with lab reference strain and clinical strains were identified, with extensive granuloma formation in the lungs of the clinical isolates (**Figure 2C**). In addition, areas of dense nuclei, representing lesion areas (highlighted in red), were observed, demonstrating clear strain- and dose-dependent effects on lung pathology (**Figure 2D**). H&E staining revealed granuloma-like lesions with macrophage infiltration in all infected groups, with increased lesion severity across clinical strains (**Figure 2E**). Ziehl-Neelsen staining confirmed numerous acid-fast bacilli within these granuloma-like structures (**Figure 2F**). No mycobacteria or granuloma-like structures were detected in uninfected controls (not shown). Together, these results demonstrate that clinical isolates generate a stronger, more persistent lung infection and pathology than the reference strain.

### 3.4 µCT supports reduction and refinement of animal use by non-invasively revealing differences between reference and clinical isolates

µCT imaging was included as a novel non-invasive readout, where we wanted to assess whether we could capture the development of *M. avium* lung pathology longitudinally, and to validate this new technique to established gold-standard readouts, namely CFU and histology. Qualitatively, a visual difference can be seen between the two inocula of the lab reference strain, with the lower inoculum still comparable to the healthy control group at endpoint, with no infiltration of inflammatory infiltrates. The arrows represent a region where air has been replaced by denser material, like these inflammatory infiltrates (**Figure 3A**). The differences between clinical isolates and lab strains captured with CFU and histology (**Figure 2**), were also confirmed by this approach. Quantitative analysis revealed a significant increase in NALV in animals infected with the clinical isolates compared with those infected with the lab strains. Among the reference strain groups, only the higher inoculum resulted in a significant increase in NALV at the endpoint, whereas infections with MYC_0069 and MYC_0068 led to significant increases as early five and seven weeks post-infection, respectively (**Figure 3B**). This distinction between the two clinical strains was not evident from CFU enumeration alone.

**Figure 3.**
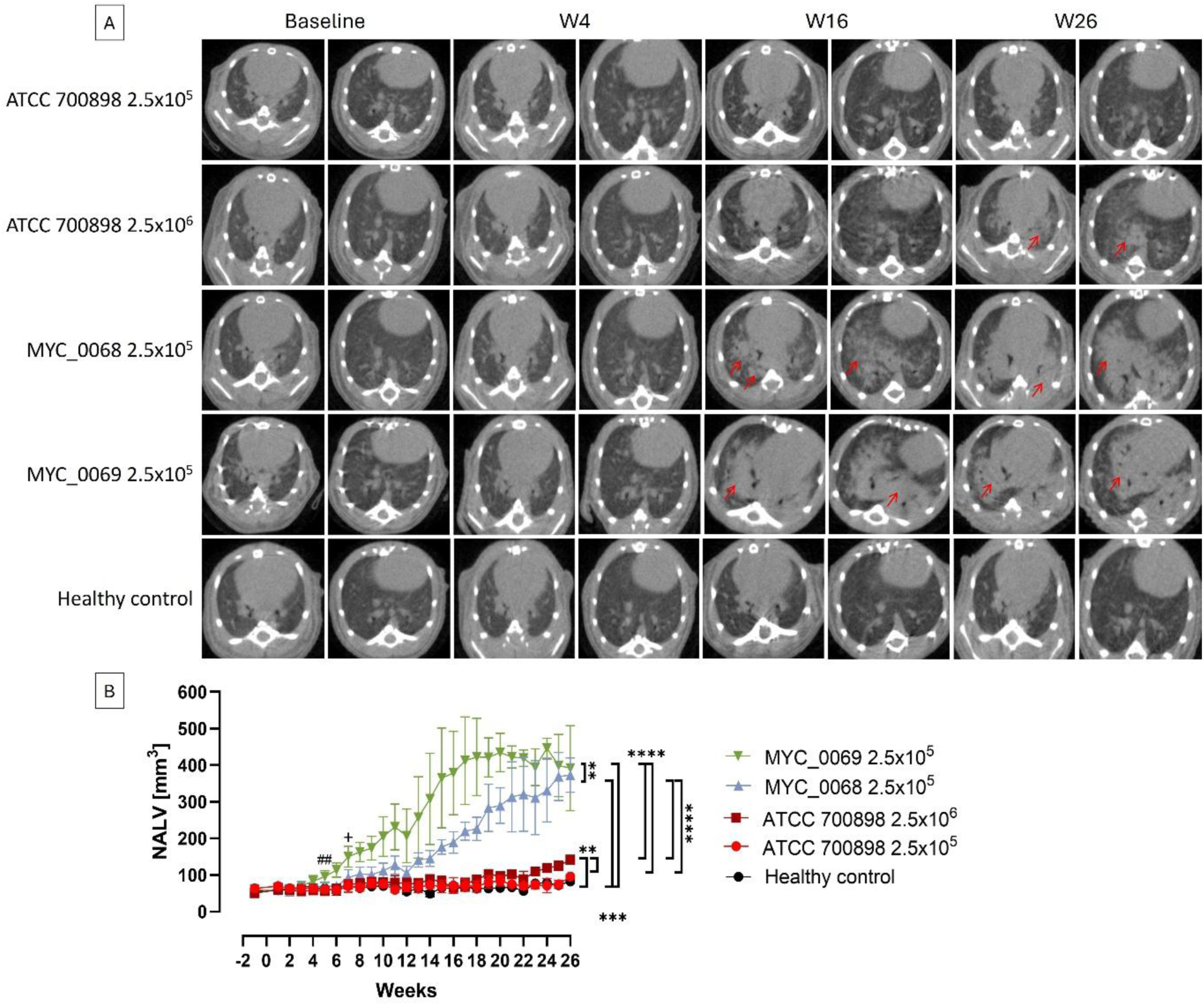
µCT allows for non-invasive assessment of progressive pulmonary pathology in lungs, while allowing differentiations between strains. **A)** Representative images of a µCT scans from mice infected with different *M. avium* strains and inoculum sizes at baseline, weeks 4, 16, and 26. Red arrows indicate a region where air has been replaced by denser material, like inflammatory infiltrates. **B)** Longitudinal quantification of lung lesion burden expressed as NALV over a 26-week follow-up period. Data are presented as mean ± SD. Group sizes were n = 8 infected and n = 4 healthy controls until week 4; n = 7 infected and n = 2 healthy until week 10; n = 6 infected and n = 2 healthy until week 16; and n = 5 infected and n = 1 healthy at week 26. “#” on the graph refers to differences at 5 weeks post infection between MYC_0069 and healthy control, “+” between MYC_0068 and healthy group. Statistical significance: *P < 0.5, **P < 0.01, ***P < 0.001, ****P < 0.0001.

Importantly, although increasing the inoculum of the laboratory strain resulted in a higher bacterial burden and detectable changes in NALV, the higher inoculum did not compensate for the intrinsically reduced virulence of the laboratory strain compared to the clinical isolates. In other words, pathology progression remained less pronounced and delayed, even under higher infectious pressure.

These observations show that non-invasive µCT readouts provide a more sensitive measure of *M. avium* lung pathology progression than conventional microbiological and endpoint-based readouts. µCT enables longitudinal assessment in the same living animal, allowing the establishment of individual baseline measurements and the monitoring of disease progression over time. This approach, which requires the use of selected clinical strains, reduces animal use while providing richer and more dynamic information than the current gold-standard endpoint analyses. Based on these findings, MYC_0069 was selected as the most relevant and virulent clinical isolate for further investigation in downstream experiments, enabling a more focused comparison of antibiotic treatment responses with the laboratory reference strain.

### 3.5 Antibiotic responses differ between reference and clinical strain infection in *Galleria mellonella*

As a next step, we aimed to determine whether treatment responses differed between the most virulent clinical isolate and the laboratory reference strain under antibiotic pressure. To this end, larvae were infected with either *M. avium* ATCC700898 or MYC_0069 and treated daily from one day until day 7 post infection with different doses of clarithromycin, allowing a direct comparison of treatment responses between infections with both strains.

Consistent with the findings reported in **Figure 1**, no evident bacterial replication was observed for the ATCC 700898 laboratory strain, and therefore no significant differences in bacterial burden were detected between treatment groups, except at the highest clarithromycin dose (100 mg/kg) at endpoint (**Figure 4A**). In contrast, with clinical isolate MYC_0069, bacterial replication was observed over time, and significantly reduced bacterial burden with clarithromycin 100 mg/kg at day 3 post infection (**Figure 4B**). As MIC values for clarithromycin, rifampicin, and ethambutol were comparable across ATCC 700898, MYC_0068, and MYC_0069 (**Supplementary table 1**), these differences in treatment response are unlikely to be explained by baseline antimicrobial susceptibility and instead appear to reflect differences in bacterial growth dynamics *in vivo*.

**Figure 4.**
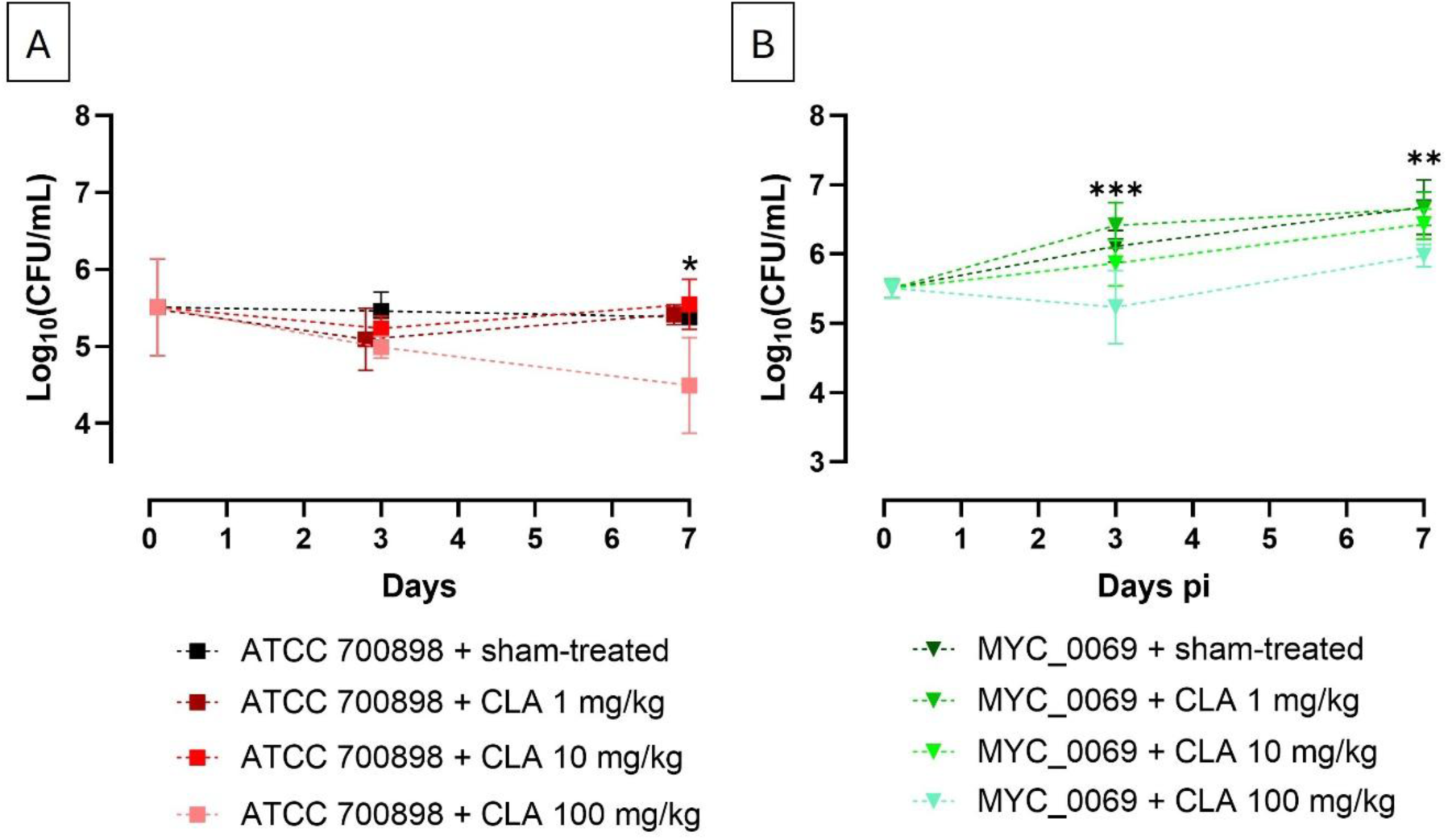
Clinical and reference *M. avium* strains show distinct clarithromycin responses in *Galleria mellonella*. **A)** Larvae were infected with 10^6^ CFU/10 µL of *M. avium* ATCC 700898. **B)** Larvae were infected with 10^6^ CFU/10 µL of *M. avium* MYC_0069. Bacterial burden (log_10_ CFU/mL) was quantified at 1 hour, 3 days, and 7 days post infection. Clarithromycin (CLA) or vehicle (0.6% acetic acid in PBS) treatment was initiated one day post infection and administered daily until day 7. Bars indicate ± SD (n = 4 per timepoint). “*” on the graph refers to differences between ATCC 700898 or MYC_0069 treated with clarithromycin 100 mg/kg. Statistical significance: **P < 0.01, ns = non-significant.

### 3.6 Classical endpoints confirm therapeutic efficacy in mice infected with a clinical strain

Treatment response was further evaluated in BALB/c mice infected with clinical isolate MYC_0069, providing a more clinically relevant context and enabling assessment in a more complex *in vivo* model. The inoculum used for this experiment was similar to the inoculum used in the experiment described in section 3.3 and 3.4, showing reproducibility between experiments (**Supplementary figure 3**).

Treatment efficacy was first evaluated using CFU enumeration. Clarithromycin monotherapy and triple-combination therapy were initiated four weeks after infection and administered daily for 12 weeks. Both regimens resulted in a significant reduction in bacterial burden, with CFU counts decreasing by approximately 2 log_10_ by week 10. At endpoint (week 16), complete bacterial eradication was observed in all mice treated with clarithromycin monotherapy (4/4) and in half of the mice receiving triple-combination therapy (2/4) (**Figure 5A**). Lung weights were recorded concomitantly with the CFU analysis, and remained stable in both treated groups, whereas a progressive increase was observed in vehicle-treated mice, indicative of ongoing disease (**Figure 5B**). Similar trends in bacterial burden and organ weight were confirmed in the spleen, supporting a systemic treatment effect (**Supplementary figure 5A–D**).

**Figure 5.**
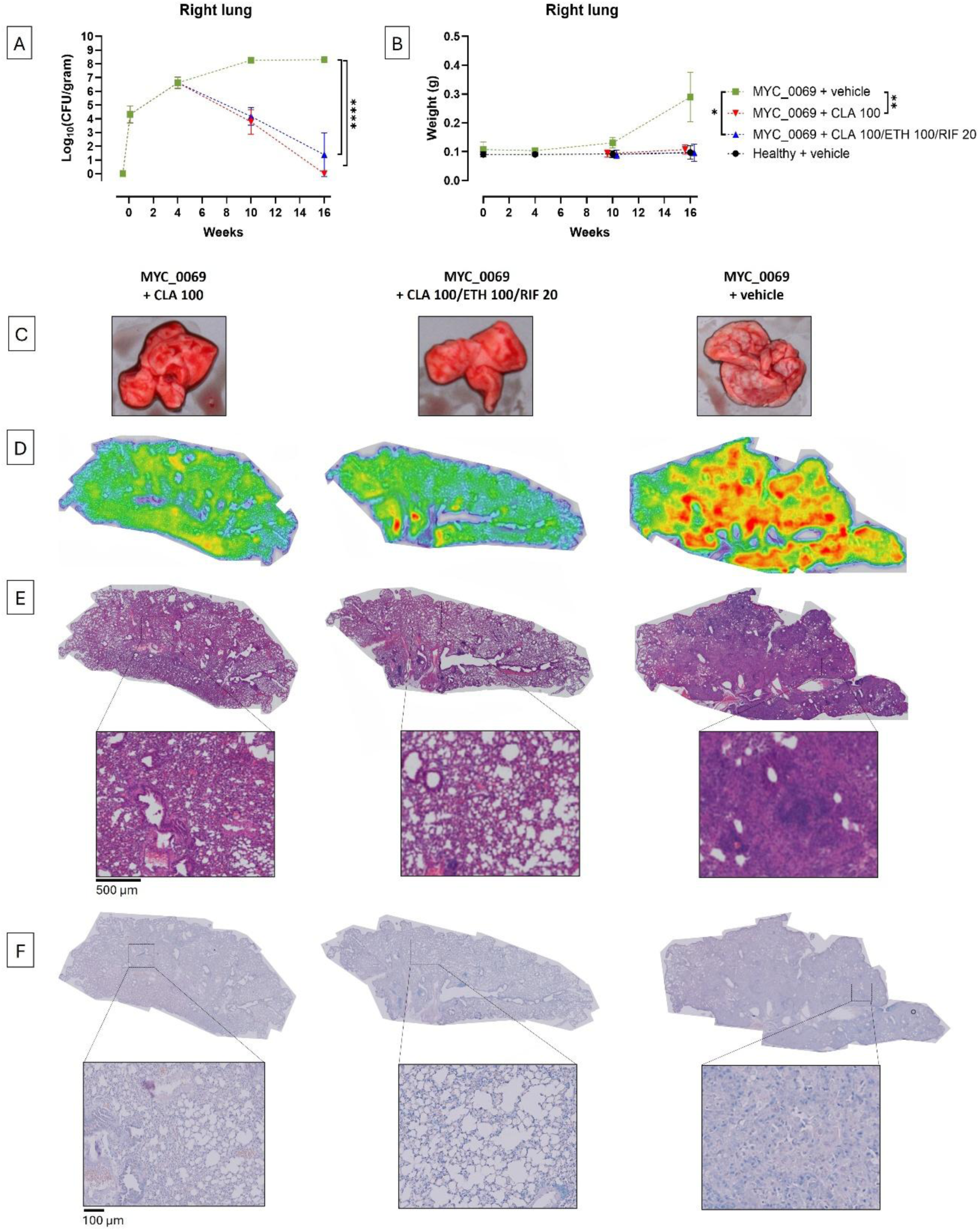
Therapeutic efficacy of clarithromycin monotherapy and standard-of-care treatment regimen in BALB/c mice infected with clinical isolate MYC_0069 can be detected with classic endpoint analysis. BALB/c mice were intranasally infected with 2.5×10^6^ CFU of *M. avium* MYC_0069. Clarithromycin (CLA) monotherapy or triple-combination therapy (CLA + ethambutol (ETH) + rifampicin (RIF)) was initiated four weeks post infection and administered daily until week 16. Vehicle-treated controls received 0.6% acetic acid in PBS. **A)** Lung bacterial burden over time, expressed as log_10_(CFU/gram), measured at day 1 and weeks 4, 10, and 16 post infection. **B)** Lung weights measured at the corresponding timepoints. Bars indicate mean ± SD (n = 4 mice per group per timepoint). At endpoint, one mouse from the healthy control group was excluded due to unintended infection with the mycobacterial strain (n = 3). **C)** Representative macroscopic images of the right lungs collected at week 16; intermediate timepoints are not shown. **D)** Heat map representations generated from H&E-stained lung sections. Red represents lesion areas, green indicates thickened parenchyma, and blue indicates uninvolved parenchymal tissue. **E)** H&E staining of lung sections showing whole-lung overviews and magnified regions (scale bar: 500 µm). **F)** Ziehl-Neelsen staining of the same lung sections shown in panel E (scale bar: 100 µm). Statistical significance: *P < 0.05, **P < 0.01, ****P < 0.0001.

To assess the impact of mono- and triple-combination therapy on lung structure, lungs were imaged at each timepoint and subjected to histopathological analysis. A clear distinction was observed between both treatment groups and the vehicle-treated group, whereas minor differences were observed between the mono- and combination therapy groups (**Figures 5C** and **5D)**. Macroscopically, lungs from vehicle-treated animals displayed visible development of granuloma-like structures, whereas lungs from both treated groups retained a largely normal appearance and lack obvious lesion formation (**Figure 5C**). These differences were also reflected in the heat map analysis, where extensive lesion-associated areas (red) were evident throughout the lungs of vehicle-treated mice, while only limited lesions were present in the clarithromycin- and triple-combination-treated groups. A larger proportion of uninvolved parenchyma (blue) was preserved in treated animals compared to vehicle controls (**Figure 5D**). Endpoint H&E staining revealed granuloma-like lesions with pronounced macrophage infiltration in the vehicle-treated group, while both treated groups showed reduced but still apparent inflammatory infiltrates (**Figure 5E**). Ziehl–Neelsen staining confirmed the presence of mycobacteria in infected groups, with a reduced bacterial presence in treated animals compared to the vehicle control. No mycobacteria or granuloma-like structures were detected in uninfected controls (not shown) (**Figure 5F**).

### 3.7 Lung µCT detects treatment efficacy in BALB/c mice infected with clinical *M. avium* strain within one week after treatment initiation, reducing dependence on terminal endpoints

To determine whether µCT imaging could non-invasively detect treatment efficacy and corroborate classical endpoint analyses, µCT was included as an additional longitudinal readout. Mice infected with the clinical isolate *M. avium* MYC_0069 were weekly imaged throughout treatment with the standard-of-care regimen or vehicle.

Qualitatively, longitudinal µCT images revealed clear differences, indicated with red arrows, between treated and vehicle-treated groups that were consistent with results of CFU and histology (see **Figure 5**). In both therapy groups, lung architecture at later timepoints closely resembled baseline scans obtained prior to infection, whereas infected, vehicle-treated mice developed extensive pulmonary pathology by endpoint (**Figure 6A**).

**Figure 6.**
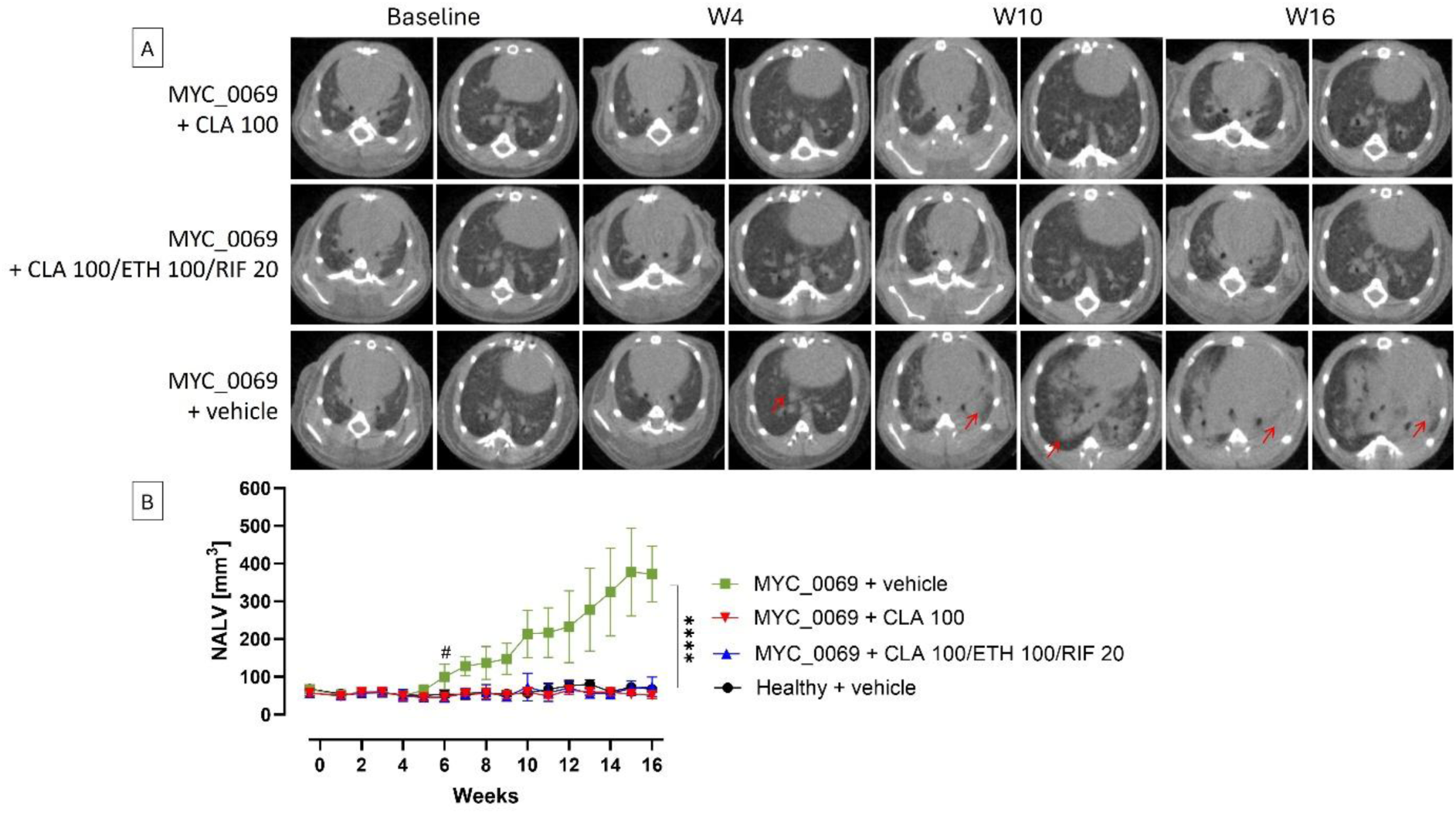
Longitudinal imaging sensitively captures early therapeutic effects in pulmonary *Mycobacterium avium* infection. **A)** Representative longitudinal µCT lung images illustrating treatment-associated changes in pulmonary pathology at week 16 post infection. Red arrows indicate a region where air has been replaced by denser material, like inflammatory infiltrates. **B)** Longitudinal quantification of lung lesion burden expressed as NALV over a 16-week follow-up period. Data are presented as mean ± SD. Group sizes were n = 32 infected and n = 16 healthy controls until day 1; n = 28 infected and n = 12 healthy until week 4; n = 8 infected + clarithromycin (CLA, 100 mg/kg), n = 8 infected + triple therapy (CLA + ethambutol (ETH, 100 mg/kg) + rifampicin (RIF, 20 mg/kg)), n = 8 infected + vehicle (0.6% acetic acid in PBS), and n = 8 healthy until week 10; n = 4 infected + CLA, n = 4 infected + triply therapy, n = 4 infected + vehicle, and n = 3 healthy at week 16. One healthy mouse was excluded at endpoint due to unintended cross-infection. “#” on the graph refers to differences at 6 weeks post infection between infected vehicle-treated group and the healthy vehicle-treated group. Statistical significance: *P < 0.05, ****P < 0.0001.

Quantitative analysis of NALV supported these observations. A statistically significant difference between treated and vehicle-treated groups was detectable as early as two weeks after treatment initiation. Both mono- and triple-combination therapy resulted in a marked and significant reduction in NALV, with no significant difference in efficacy between the two treatment regimens (**Figure 6B**).

Together, these findings demonstrate that µCT imaging sensitively captures treatment-induced changes in pulmonary pathology and confirms therapeutic efficacy detected by classical readouts, while enabling early, longitudinal assessment within the same animals with a reduced need for invasive endpoints.

## 4. Discussion

In this study, we demonstrate that clinically derived *M. avium* isolates exhibit markedly different virulence and disease dynamics compared to a commonly used laboratory reference strain, and that these differences are consistently captured across human macrophage*, Galleria mellonella,* and µCT-compatible mouse infection models. Importantly, we show that longitudinal µCT imaging enables early, non-invasive detection of treatment-associated changes in pulmonary pathology, confirming and extending classical endpoint readouts while reducing reliance on invasive sampling and uniquely enabling individual baseline measurements for powerful therapeutic efficacy assessment.

Across experimental systems, recent clinical isolates displayed enhanced bacterial growth and pathology relative to the reference strain. Although extracellular growth kinetics were comparable *in vitro*, clinical isolates clearly replicated more intracellularly in THP-1-derived macrophages, indicating strain-dependent differences in host-pathogen interactions, key to mycobacterial infections. Similar trends were observed across the *in vivo* models, where clinical isolates established higher and more persistent bacterial burdens and induced more pronounced pulmonary pathology. Comparable observations have been reported for *Mycobacterium tuberculosis*, where laboratory-adapted strains fail to fully recapitulate clinically relevant infection phenotypes and therapeutic responses (23,24). Collectively, these findings underscore the importance of strain selection in preclinical *M. avium* research and highlight the limitations of relying solely on long-established laboratory strains.

Among our collection of recent *M. avium* strains isolated from patients with MAC pulmonary disease, MYC_0069 displayed a particular growth phenotype, characterized by robust growth both intracellularly and *in vivo*, supporting the selection of this strain for downstream therapeutic evaluation. Notably, these observations indicate that strain selection not only influences disease severity but also infection dynamics. While bacterial burdens reported for ATCC 700898 were comparable to those observed in this study during the first four weeks following infection, the reference strain subsequently plateaued at ∼7 log_10_ CFU, whereas MYC_0069 continued to replicate and reached ∼8.3 log_10_ CFU before stabilizing (25). In line with these findings, previous studies using the same reference strain also reported minimal bacterial replication throughout the course of infection (22). These findings suggest a more progressive *in vivo* infection phenotype for the clinical isolate, providing a broader dynamic range for evaluating treatment efficacy. The apparent adaptation of this isolate to the *in vivo* environment may also enhance the translational relevance of the model.

The marked differences in bacterial replication observed in *G. mellonella* further highlight important implications for therapeutic evaluation. Whereas the ATCC 700898 reference strain showed limited replication and apparent bactericidal responses at the highest treatment dose, MYC_0069 remained actively replicative throughout infection, resulting primarily in dose-dependent growth inhibition. This suggests that laboratory strains may overestimate compound efficacy, while actively replicating clinical isolates provide a more stringent and clinically relevant platform for preclinical drug assessment.

While the choice of infecting strain is critical for establishing a clinically relevant disease model, sensitive and ethically responsible tools are equally needed to capture disease progression and therapeutic responses over time. A central finding of this study is the added value of longitudinal µCT imaging as a sensitive, non-invasive readout of disease progression and treatment response. µCT-derived NALV closely reflected strain-dependent differences in progression of lung pathology, as well as differences in bacterial burden, while offering longitudinal insight into disease development that cannot be obtained from endpoint-based readouts such as CFU enumeration and histopathology alone. Notably, treatment-associated reductions in lung pathology were detectable by µCT within two weeks of therapy initiation, demonstrating the ability of longitudinal imaging to capture early therapeutic effects in real time before final endpoint assessment. The ability to repeatedly monitor the same animals throughout infection and treatment additionally enables the detection of relapse or treatment failure and supports reduction and refinement of animal use in accordance with the 3Rs. These findings further expand previous applications of µCT in pulmonary infection and disease models (17). In models relying on laboratory strains with limited disease progression, treatment-associated changes may be too subtle to detect. This highlights the value of active infection models with clinically relevant isolates, in which longitudinal imaging can provide additional insights into the dynamics of the host response.

Having established MYC_0069 as a stringent infection model, therapeutic evaluation confirmed robust efficacy of both clarithromycin monotherapy and triple-combination therapy. Consistent with previous murine studies, showing no differences between treatment regimens with respect to bacterial clearance kinetics, endpoint bacterial burden, or pulmonary pathology (22,25), and together with reports of substantial variability in pathology among clinical *M. avium* isolates (26), our findings support both the robustness of clarithromycin-mediated treatment responses and the importance of using clinically relevant isolates for disease modeling.

The comparable efficacy observed between clarithromycin monotherapy and triple-combination therapy should nevertheless be interpreted with caution. Although both regimens achieved similar reduction in bacterial burden in this model, clarithromycin monotherapy is not recommended in clinical practice because macrolide resistance can emerge during treatment, compromising future treatment options. The current standard-of-care remains multidrug therapy for *M. avium* disease. Importantly, similar findings have previously been reported in murine models of chronic *M. avium* infection, in which clarithromycin monotherapy achieved comparable efficacy to multidrug therapy, without evidence of resistance emergence (22). Our findings further support the central role of clarithromycin as the primary driver of activity within current MAC treatment regimens and raise broader questions regarding the contribution of rifampicin to treatment efficacy. Consistent with this interpretation, It was recently demonstrated that omadacycline-containing regimens achieved efficacy comparable to rifampicin-containing regimens (25). Although caution is warranted when extrapolating preclinical findings to patients, our results emphasize the continued need to identify more effective companion drugs that can provide additional benefit beyond that provided by clarithromycin alone.

No clear indication of acquired resistance was observed during the 12-week treatment period, including the clarithromycin monotherapy group, where resistance development might theoretically be expected to occur more readily than under combination therapy. This is supported by the continuous decline in bacterial burden throughout treatment and the achievement of complete bacterial eradication at the experimental endpoint in all mice receiving monotherapy. Yet, the presence of a small number of residual persister bacteria below the limit of detection cannot be excluded. As no viable bacteria were recovered in these groups, MIC determination and whole-genome sequencing could not be performed on endpoint isolates, precluding definitive confirmation or exclusion of acquired resistance.

This study has limitations that should be acknowledged. Treatment experiments were performed using a single clinical isolate, and future studies incorporating multiple clinically diverse strains will be important to generalize these findings. In addition, although longitudinal CFU measurements clearly demonstrated progressive expansion of the clinical isolates over time, inclusion of an early post infection timepoint would have enabled a more precise assessment of initial bacterial establishment and subsequent *in vivo* growth kinetics. Additionally, cavitary disease was not observed in this model, which would likely alter drug penetration. Pharmacokinetic assessments were beyond the scope of this work.

In conclusion, this study demonstrates that clinically relevant *M. avium* isolates reveal disease dynamics that are not captured by laboratory reference strains and that more closely resemble the pathology observed in human disease. Longitudinal µCT imaging provides a powerful approach for sensitively assessing both disease progression and treatment efficacy. Together, these findings establish a clinically relevant chronic mouse infection model based on a virulent clinical isolate that is well suited for the preclinical evaluation of novel anti-MAC drug candidates. By combining clinically relevant bacterial strains with non-invasive longitudinal imaging, this approach has the potential to improve the translational predictability of preclinical drug evaluation while reducing experimental timelines, resource requirements, and animal use.

## 5. Supplementary figures

**Supplementary figure 1.**
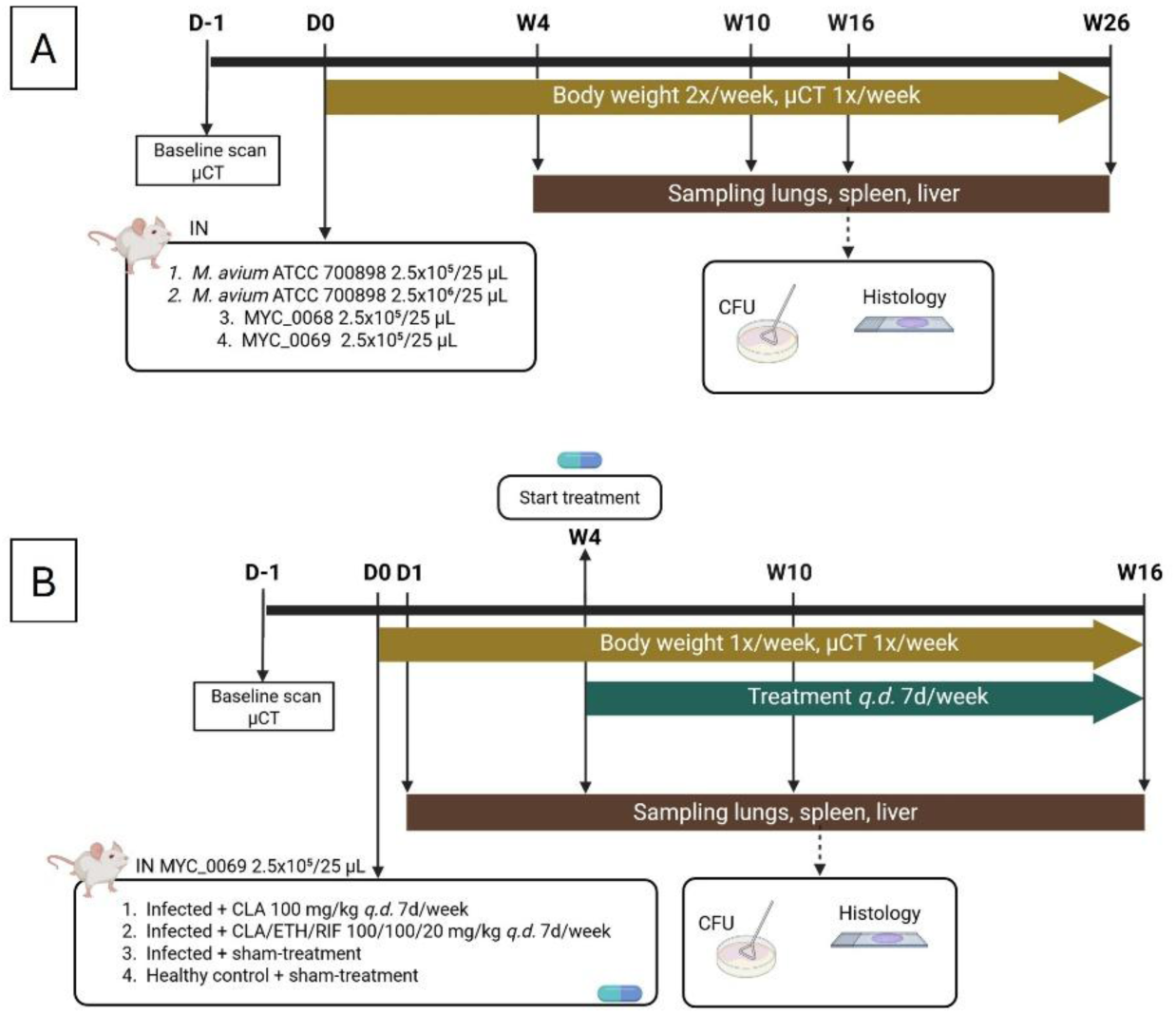
Experimental outline. **A)** Mouse experiment 1. **B)** Mouse experiment 2.

**Supplementary figure 2.**
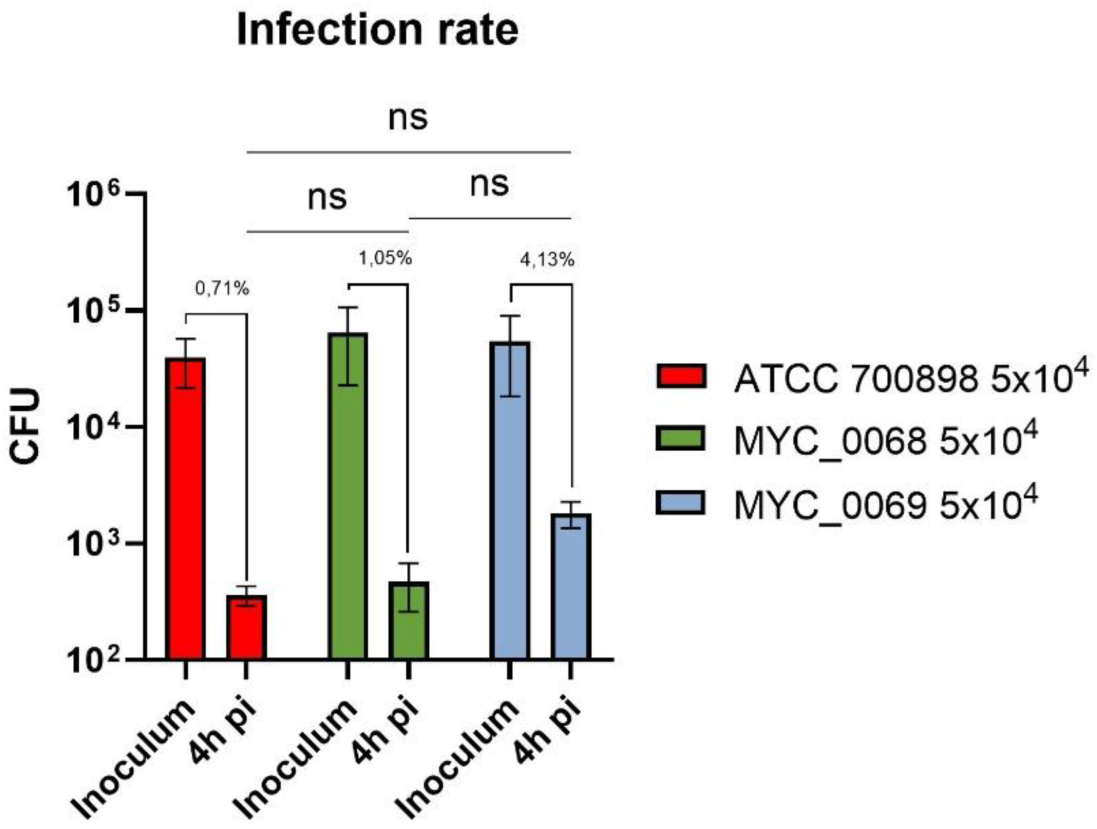
*In vitro* infection rate differs between reference and clinical *M. avium* strains. Macrophage infection rates for each *M. avium* strain, expressed as the proportion of infected cells. Data are presented as mean ± SD from biological replicates. Statistical significance: ns = non-significant.

**Supplementary figure 3.**
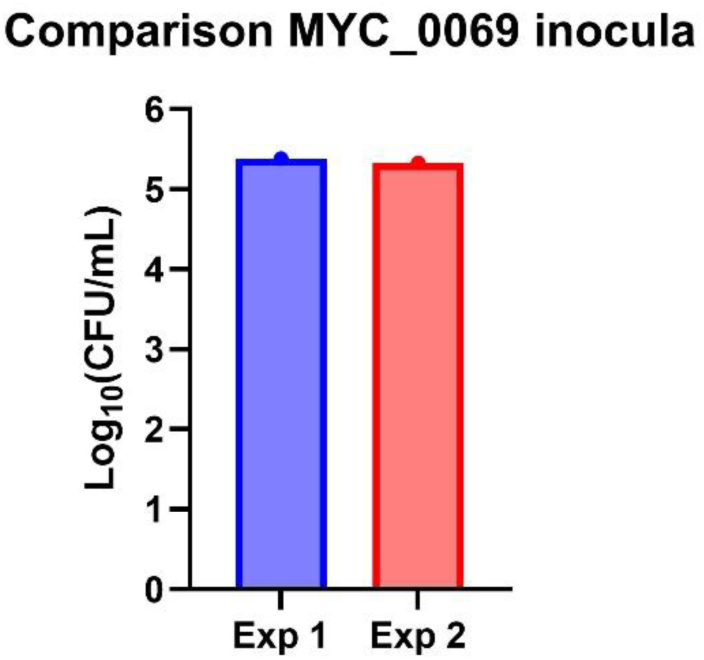
Comparable inoculum levels of the clinical *M. avium* isolate MYC_0069 across murine experiments. Bacterial load of the *M. avium* clinical isolate MYC_0069 used for inoculation in two independent murine experiments.

**Supplementary figure 4.**
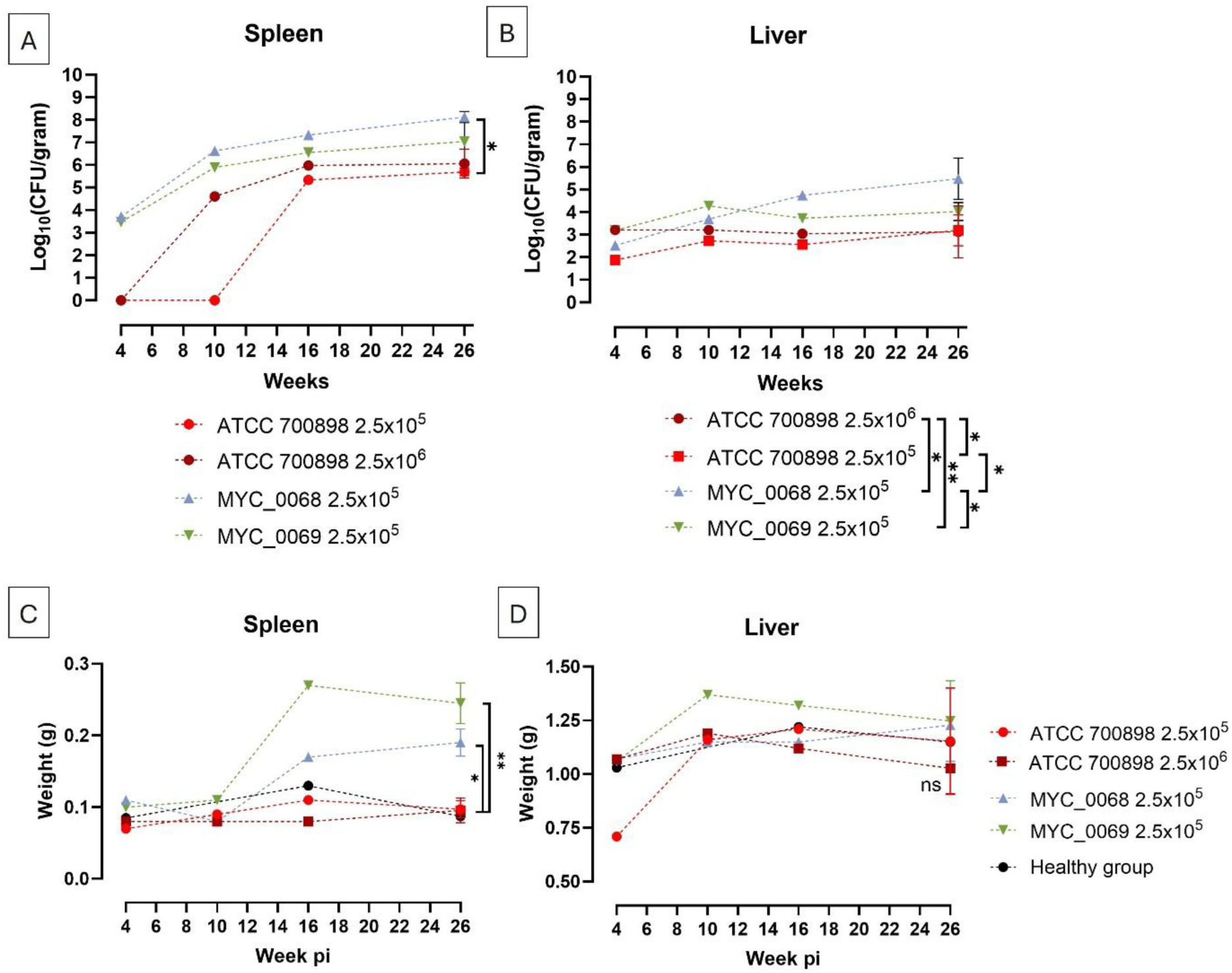
Clinical *M. avium* isolate exhibits increased extrapulmonary bacterial burden compared with the reference strain. **A)** Spleen bacterial burden over time. BALB/c mice were infected with 2.5×10^6^ CFU of *M. avium* ATCC 700898 or 2.5×10^5^ CFU of *M. avium* ATCC700898, MYC_0068, or MYC_0069 at indicated timepoints post infection. **B)** Liver bacterial burden in the liver of the same animals at corresponding timepoints. **C)** Spleen weights at corresponding timepoints. **D)** Liver weights at corresponding timepoints. Data are represented as mean ± SD for each group (n = 1/2 per infected/healthy group at week 4, n = 1 per infected group at week 10, n = 1 per group at week 16; n = 5/1 per infected/healthy group at week 26). Statistical significance: *P < 0.05, **P < 0.01, ns = non-significant.

**Supplementary figure 5.**
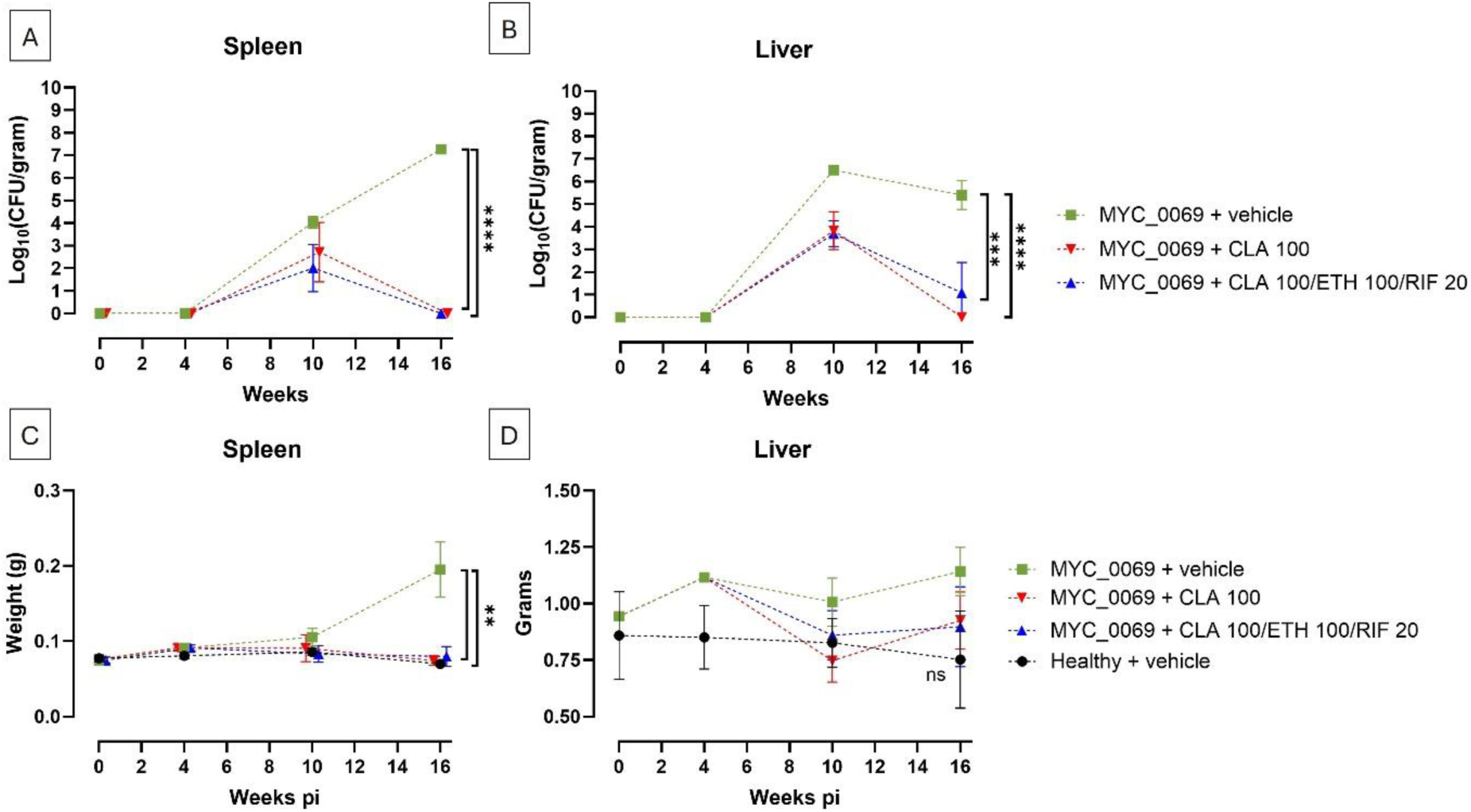
Treatment reduces extrapulmonary bacterial burden in mice infected with the clinical *M. avium* isolate MYC_0069. **A)** Spleen bacterial burden over time. BALB/c mice were intranasally infected with 2.5×10^6^ CFU of MYC_0069, and spleen bacterial burden was quantified as log_10_(CFU/gram) at day 1 and weeks 4, 10 and 16 post infection. Clarithromycin (CLA) monotherapy or triple-combination therapy (CLA + ethambutol (ETH) + rifampicin (RIF)) was initiated four weeks post infection and administered daily until week 16. Vehicle-treated controls received 0.6% acetic acid in PBS. **B)** Liver bacterial burden of the same animal at corresponding timepoints. **C)** Spleen weights measured at corresponding timepoints. **D)** Liver weights measured at corresponding timepoints. Bars indicate mean ± SD (n = 4 mice per group per timepoint). At endpoint, one mouse from the healthy control group was excluded due to unintended infection with the mycobacterial strain (n = 3). Statistical significance: **P < 0.01, ****P < 0.0001, ns = non-significant.

**Supplementary table 1.** Minimal Inhibitory Concentration (MIC) comparison between *M. avium* strains. The values shown represent the average of three determinations.

| Minimal Inhibitory Concentration (mg/L) |  |  |  |
| --- | --- | --- | --- |
| *M. avium* strain | Clarithromycin | Rifampicin | Ethambutol |
| ATCC 700898 | 0.88 | 0.28 | 5 |
| MYC_0068 | 4 | 0.57 | 32 |
| MYC_0069 | 1.92 | 0.22 | 8.67 |

## 7. Author contributions

Conceptualization: T.V.W., W.B., A.R.S., G.J.W., N.L., G.V.V., E.A.; data curation: T.V.W., W.B., E.D.P.; formal analysis: T.V.W.; investigation: T.V.W., W.B.; methodology: T.V.W., W.B., K.B.; supervision: A.R.S., G.J.W., G.V.V., E.A.; writing – original draft: T.V.W.; writing – review and editing: T.V.W., W.B., A.R.S., G.J.W., E.D.P., N.L., K.B., G.V.V., E.A.; visualization: T.V.W. All authors have read and approved the final version of the manuscript.

## 9. Acknowledgments

We sincerely thank all colleagues who contributed to this work through valuable discussions, technical support, and scientific expertise.

